# jsPCA: fast, scalable, and interpretable identification of spatial domains and variable genes across multi-slice and multi-sample spatial transcriptomics data

**DOI:** 10.1101/2025.09.16.676466

**Authors:** Ines Assali, Paul Escande, Franck Picard, Paul Villoutreix

**Affiliations:** Aix-Marseille Université, MMG, Inserm U1251, Turing Centre for Living systems, Marseille, France; Institut de Mathématiques de Toulouse; UMR 5219, Université de Toulouse, CNRS; UPS, F-31062 Toulouse Cedex 9, France; Université de Lyon, ENS de Lyon, Université Claude Bernard, CNRS UMR 5239, INSERM U1210, Lyon, France

**Keywords:** Spatial transcriptomics, Spatial statistics, Machine learning, Spatial domains, Spatially variable genes

## Abstract

Spatial transcriptomics technologies record genome-wide measurements of gene expression with high spatial resolution. These technologies generate large and high-dimensional datasets requiring efficient automated methods for their analysis. We introduce joint spatial PCA (jsPCA), a novel, fast, scalable and interpretable method for the automatic identification of spatial domains and variable genes in multi-slice and multi-sample spatial transcriptomics data. jsPCA relies on a simple mathematical formulation of a spatial covariance defined as the product of the gene expression covariance with the spatial autocorrelation. The principal components of this spatial covariance yield a biologically meaningful low-dimensional representation. From this representation, spatial domains are derived by simple clustering and spatially variable genes are identified directly from the principal component coefficients. A joint representation of multiple slices and samples without spatial alignment is obtained by computing common principal components via joint diagonalization. By leveraging data sparsity and non-convex manifold optimization, jsPCA leads to computing time in the order of seconds to minutes, substantially outperforming state-of-the-art approaches. We benchmarked jsPCA against 10 state-of-the-art methods on two reference databases. Our approach demonstrated excellent performance, comparable or better than state-of-the-art methods, while being much faster, interpretable, and scalable to very large datasets.

## Introduction

Spatial transcriptomics (ST) has changed the way we understand biological systems by offering a perspective that combines gene expression and spatial information within tissues. Unlike traditional single-cell sequencing technologies that dissociate cells from their microenvironment, ST preserves the spatial structure of tissues while capturing gene expression profiles, enabling an in-depth analysis of cellular expression *in situ* [1]. This technological advance enables not only to map the location of gene expression in tissues but also to understand the mechanism of cell-cell interaction [2], to identify cell types according to their localization and molecular profile [3], and to better understand a variety of biological processes, including tissue pathology [4], immune responses [5], and embryonic development [6]. These advances open up new possibilities for personalized medicine and the development of targeted treatments [7]. However, this technology faces significant challenges in terms of data sparsity, very high dimensionality, and technical and biological noise [8]. These fundamental data limitations require innovative computational strategies capable of efficiently normalizing, reducing dimensionality while preserving essential biological insights.

Specifically, the task of identifying spatial domains that are both consistent in terms of expression and spatial organization has gathered attention in the past few years [9]. Several approaches have been proposed to solve this task as a problem of spatially aware clustering. Some of these methods rely on graph neural networks, such as STAGATE, which is a graph attention auto-encoder framework [10], SpaceFlow, which is a deep learning framework based on spatially regularized deep graph networks [11], and CCST (Cell Clustering for Spatial Transcriptomics), which is an unsupervised cell clustering method based on a graphical neural network [12]. Another set of methods relies on a Bayesian framework, such as BASS which is a multi-scale, multi-sample approach to spatial transcriptomic analysis [13]. BASS explores data at different scales by simultaneously clustering cell types and detecting spatial domains. These analyses are integrated into a hierarchical Bayesian modeling framework. Similarly, BayesSpace is a Bayesian statistical method that uses the neighborhood information [14]. Finally, some approaches have been proposed to extend the classical Principal Component Analysis approach to spatial transcriptomics, such as SpatialPCA [15], which is based on probabilistic PCA, it includes location information as an additional input and uses a kernel matrix to model spatial correlations between tissue sites. Similarly, GraphPCA is a dimension reduction algorithm that integrates a graph based regularization of principal component analysis (PCA) to improve the analysis of spatial transcriptomic data [16]. These methods are complex, not always interpretable, or require tuning several hyperparameters, and can be slow in their execution.

In addition to spatial domain identification, an important step in spatial transcriptomics analysis is identifying which genes are spatially variable (SVGs) [17]. These genes exhibit non-random expression patterns across tissue, often reflecting specific biological functions or distinct cellular microenvironments. Their identification not only highlights relevant spatial patterns, but can also be used to reduce the dimensionality of the data. SVGs are similar to the selection of highly variable genes (HVGs) in single-cell transcriptomics; however, they differ in that they preserve the spatial relationships specific to biological samples. SVGs thus play a central role in many downstream analyses, such as the exploration of pathological mechanisms [17]. Numerous approaches have been developed for this task, ranging from methods based on Gaussian processes (SpatialDE) [18] to mixed or non-parametric models (SPARK [19], SPARK-X [20]), including self-organizing maps (SOMDE) [21], statistical enrichment in spatial networks (Giotto) [22], Gaussian processes based on nearest neighbors (nnSVG) [23], and spatial autocorrelation measures such as Moran’s I (implemented in Seurat [24], among others These approaches have mostly been developed independently of domain identification methods and are usually used as a preprocessing step to reduce the dimensionality of the data. Moreover, they rely on sometimes complicated statistical models.

In this study, we propose joint spatial Principal Component Analysis (jsPCA), a novel computational method for identifying spatial domains and spatially variable genes from spatial transcriptomics data that addresses the limitations of existing methods. jsPCA generalizes sPCA, an established method for population genetics previously applied to genetic variability using allelic frequency data and spatial coordinates [25]. We previously introduced an initial formulation of jsPCA to study single-cell morphometrics data within biological tissues obtained by microscopy imaging [26]. Adapting jsPCA to spatial transcriptomics data generates new methodological challenges related to the high dimensionality of this type of data. We solved these by exploiting the sparse nature of the matrices and non-convex optimization on manifold techniques [27, 28]. jsPCA efficiently integrates spatial information with gene expression profiles to produce a low-dimensional representation of the data that enable the identification of spatial domains and spatially variable genes. jsPCA is well adapted for multi-slice and multi-sample data.

## Methods

### Data preprocessing

To optimize the detection of biological signals, we carried out a three steps pre-processing of the gene expression data using the Scanpy library [29]. Firstly, we applied a prevalence-based selection criterion, selecting only genes expressed in at least 20 cells/spots. This eliminates potentially artefactual and poorly informative variables. Next, we transformed the data using Pearson’s residual normalization, which is a technique that scales expression levels based on the relationship between mean and variance, allowing us to correct any technical biases while preserving biologically significant variability. Then we normalized the data, transforming each gene expression profile into z-scored values, guaranteeing zero mean and unit variance. This transformation makes expression levels across different genes comparable, regardless of their initial expression amplitude, facilitating the robust identification of relevant biological patterns. Normalization is performed separately for each slice and sample.

### Construction of spatial neighborhood graph

To encode the spatial information contained in ST, we built a neighborhood network based on the spatial coordinates of the spots/cells. For this, we applied the *k*-nearest neighbor (*k*-NN) algorithm based on Euclidean distance, adapting the classic elbow method [30] to determine the optimal number of *k* neighbors. This modified version selects *k* by analyzing the variation in average distances to the *k*-th neighbors and identifying the point where the slope of the distance curve changes most strongly. The connectivity of the resulting network is encoded into a sparse adjacency matrix *C*. The rows and columns of *C* correspond to the spots/cells and *C*(*i, j*) = 1 if *j* ∈ *N* (*i*), *C*(*i, j*) = 0 otherwise. *N* (*i*) represents the set of neighbors of spot *i*. We denote by *L* the row-normalized version of the matrix *C*, such that the sum of each row is equal to one.

### Joint spatial principal component analysis (jsPCA)

jsPCA combines principal component analysis and spatial auto-correlation for multivariate spatial variables and multiple samples. Figure 1 shows a global overview of the method. It is based on the idea of finding an embedding of the data that represents best both the covariance, as in a classical PCA approach, and the spatial auto-correlation (also known as Moran’s index) derived from the spatial information [25, 31, 26]. In the context of spatial transcriptomics data, the problem consists in identifying gene expression patterns while preserving the spatial organization of tissues. Transcriptomic covariance is calculated using the *n* × *p* gene expression matrix *X*, and the Moran’s index is calculated as *X*^*T*^ *LX*(*X*^*T*^ *X*)^−1^.

**Figure 1.**
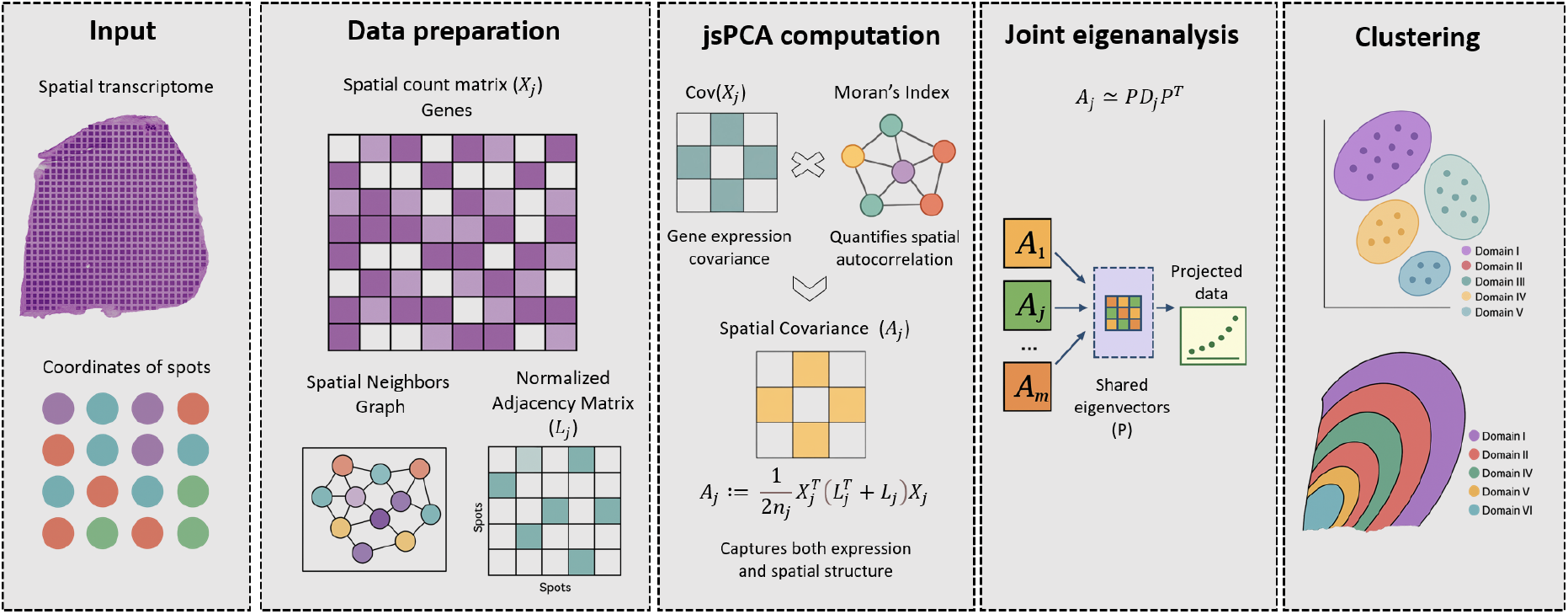
Overview of the Joint spatial PCA (jsPCA) approach. jsPCA is a joint spatial principal component analysis method that simultaneously integrates variation in gene expression and spatial structure in spatial transcriptomics (ST) data. It starts with a ST dataset consisting of a gene expression matrix X_j_ and the spatial coordinates of tissue cells/spots. From these coordinates, we construct a spatial neighborhood graph, represented by a normalized adjacency matrix L_j_, which encodes the relationships between spots. For each dataset, jsPCA then calculates a spatial covariance matrix A_j_ by combining gene expression covariance and spatial autocorrelation (measured by Moran’s index). A joint eigenvalue analysis is then performed on the set {A_1_, …, A_m_}, producing a shared basis of eigenvectors P describing the main spatio-molecular patterns. Each dataset is projected into this latent space via X_j_ P, and these representations are then used to group spots into coherent spatial domains within the tissue while the principal components can be directly interpreted in terms of spatially variable genes.

For a given sample, with expression data *X* and normalized adjacency matrix *L*, representing best both the covariance and the spatial autocorrelation can be modeled as the question of identifying the main directions of the product matrix between these two quantities: 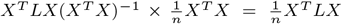. It was previously shown in [25] that the principal components (PCs) of this product matrix can be found as the eigenvectors *P* with eigenvalues *D* of the symmetrized matrix, which we call spatial covariance matrix:

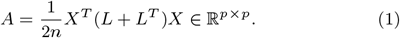

To tackle multiple datasets, such as multislice or multisample ST datasets, we need to be able to simultaneously process multiple datasets with full consideration of the spatial structure of each individual slice or sample [26]. Given a collection of samples {*X*_*j*_, *L*_*j*_}, jsPCA computes a joint diagonalization of the corresponding spatial covariance matrices, this can be summarized by finding the common set of eigenvectors *P* such that

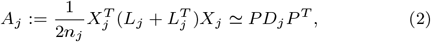

where *D*_*j*_ is the diagonal matrix corresponding to the eigenvalues specific to sample *j*. We show in the supplementary material how we obtain numerically the joint eigenvectors by non-convex optimization on a manifold because of the large size of the matrices in ST (Supplementary text 1).

The new representation 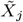 of the initial data *X*_*j*_ is obtained by projecting the data on the eigenvectors *P*. In practice we compute 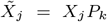 where *P*_*k*_ represents the top *k* eigenvectors of the common set of eigenvectors *P*. In the case of the analysis of a unique ST dataset, 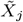 is obtained equivalently by projecting *X*_*j*_ on the eigenvectors of the spatial covariance *A*_*j*_. In the experiments below, the actual value of *k* is chosen by cross-validation in a range of 10 to 50.

Several approaches for the computation of spatial domains in spatial transcriptomics have been derived from a formulation of Principal Component Analysis with a spatial component. In the supplementary material (Supplementary text 2), we provide a formal comparison with these four methods (SpatialPCA [15], GraphPCA [16], RASP [32], nichePCA [33]). jsPCA is distinguished by its formulation, which lacks a manual weighting parameter to balance spatial versus expression data. In contrast to other methods, jsPCA has a very simple and general formulation.

### Spatial Domain Identification

For spatial domain identification, we use the low dimensional representation 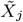 defined above, on which we apply a simple clustering algorithm, the Gaussian Mixture Model (GMM). In our experiment, the number of domains is given by the ground truth annotation, but could in principle be obtained by minimizing the Akaike Information Criterion or the Bayesian Information Criterion [34]. The labels obtained can then be reported on the tissue and compared to the ground truth.

Following clustering, we applied a refinement step, as is common in most spatial domain identification methods [9]. This refinment step was initially proposed in SPAGCN [35] and consists in a very simple step. For each spatial location, we compare its predicted label to the ones of its direct neighbors. If more than half of them have a different label, then we relabel the current one with the most common one within its direct surrounding.

### Spatially Variable Genes

Our approach naturally identifies a list of genes (the features) that explain the spatial variation within the data [36]. This list can be directly obtained from the highest contributions to the spatial principal components, a rapid and reliable method to identify important features. This approach is related to the identification of spatially variable genes (SVG) which commonly rely on a statistical testing procedure. SPARK-X for instance [20] relies on the detection of SVGs using a robust covariance test simultaneously integrating gene expression and the spatial organization of cells/spots. Specifically SPARK-X constructs a test statistic based on the trace of the product between the expression and distance covariance matrices and evaluates it with a *χ*^2^ mixture distribution to obtain calibrated p-values for selecting significant genes. jsPCA offers a simpler and more computationally efficient procedure, based on a feature contribution framework.

### Datasets and Method Evaluation

Our approach was first evaluated using the Visium 10x dataset of human dorsolateral prefrontal cortex (DLPFC) [37]. This dataset was obtained as part of the spatialLIBD project (http://spatial.libd.org/spatialLIBD). The dataset is composed of 12 slices acquired from three different individuals (four slices per sample). The number of genes per slice is 33538, with a median of 3844 spots. The samples were annotated manually, each sample comprising five or seven layers, including six cortical layers (L1 to L6) and white matter (WM), see Figure 3a.

In the first scenario, we evaluated the accuracy of spatial domain identification by jsPCA. To do so, we benchmarked on the same dataset jsPCA against a range of representative methods, including PCA-based approaches such as SpatialPCA [15], GraphPCA [16] and NichePCA [33], Bayesian methods such as BASS [13] and BayesSpace [14], graph-based methods such as STAGATE [10], CCST [12], and Spaceflow [11], as well as simpler clustering approaches including Leiden and Louvain [38]. This selection ensures a fair comparison within the same methodological family as jsPCA, while also covering a broad range of alternative approaches. Each spatial clustering method applies a specific pre-processing technique (Table S8). In this comparison, which is intended to provide a representative overview of commonly used approaches rather than an exhaustive evaluation, we adopted the optimal strategies recommended by the authors of each method. To have a fair comparison with the other methods, which do not handle multi-slice and multi-sample datasets, we performed a monoslice analysis using jsPCA, where we considered each slice of the 12 slices separately. For each slice, we projected the expression matrix onto the first *k* eigenvectors, then performed clustering in this reduced-dimension space using GMM. Instead of choosing *k* in advance, we tested a range of values. For each value, we evaluated the clustering using the silhouette score, which reflects how well the clusters are separated. We then smoothed the resulting curve with a Savitzky–Golay filter to make the trend clearer and selected *k* at the elbow point. A representative example of the resulting curve is shown in Supplementary Figure S7.

In the second scenario, we performed a multi-slice analysis with jsPCA, exploring three configurations: 2 slices, 4 slices and 12 slices. In each of these cases, we considered jointly groups of slices, for which we computed a joint embedding with jsPCA and performed clustering simultaneously. In the 2-slice analysis, we examined each pair of nearby slices, separated by a gap of 10 micrometers. In the 4-slice analysis, we studied groups of 4 slices belonging to the same sample with gaps of 10 micrometers, 300 micrometers and 10 micrometers between them. Finally, in the 12-slice analysis, we studied the three samples together, each sample consisting of 4 slices with the same gaps (10 micrometers, 300 micrometers and 10 micrometers).

We also evaluated our approach on the MOSTA (Mouse Organogenesis Spatiotemporal Transcriptomic Atlas) [39] dataset, which consists of 53 sagittal sections of mouse embryos collected at developmental stages from E9.5 to E16.5 and profiled using Stereo-seq technology. The slices contain a median of 84811 spots and 27810 detected genes, providing a detailed representation of transcriptomic organization during development. For a fair comparison, we first carried out a monoslice analysis with jsPCA, in the same manner as with the DLPFC dataset, treating each sagittal section independently, and benchmarked the results against established methods such as STAGATE, SpatialPCA, GraphPCA, and BASS, using the preprocessing strategies recommended by their authors. We then extended the analysis to a multi-slice setting by jointly considering all sections from the same developmental stage, covering E9.5 to E13.5 (5 slices at E9.5, 4 at E10.5, 4 at E11.5, 6 at E12.5, and 4 at E13.5).

Clustering performance was estimated using Adjusted Rand Index (ARI) and Normalized Mutual Information (NMI). To evaluate scalability, we measured the elapsed time and peak memory usage of our approach as well as the other approaches, on a machine with 32 GB of RAM and 24 cores (Ubuntu v.24.04), while all computations for the MOSTA dataset were executed on the IFB (Institut FranÇais de Bioinformatique) national cluster, which provides high-performance computing resources, using a node allocated with 5 CPUs and 1 TB of RAM.

## Results

To measure the ability of jsPCA at identifying spatial domains, we first evaluated it and compared it to state-of-the-art methods on the DLPFC reference dataset. We showcase the ground truth (the six manually annotated layers and white matter (WM) of the DLPFC dataset) as well as the domains identified (cluster assignments) by the different methods for sample 151673 (Fig. 2a). This comparison demonstrates the well-balanced performance of jsPCA for the identification of spatial domains in spatial transcriptomics. With an ARI of 0.62 and an NMI of 0.71, jsPCA achieves performances comparable to, or better than, the best existing approaches, such as SpatialPCA (ARI 0.57, NMI 0.70), BASS (ARI 0.59, NMI 0.70), and GraphPCA (ARI 0.47, NMI 0.65). Importantly, jsPCA is far more efficient, requiring only 22.7 seconds and 623.7 MB of memory, compared to 235.2 s / 7571.7 MB for SpatialPCA, 306.1 s / 3504.2 MB for BASS, and 624.4 s / 457.7 MB for GraphPCA. Unlike Bayespace, with its extremely high memory consumption (8699.9 MB) and longer runtime (893.7 s), or CCST, with its lower ARI and NMI scores (0.31 and 0.46) and higher memory usage (2137.2 MB), jsPCA offers a good compromise between accuracy and computational efficiency. In addition, our method outperforms Spaceflow (ARI 0.17, NMI 0.45), which achieves lower performance and produces noisier segmentation. As expected, algorithms that do not take spatial information into account, such as Leiden (ARI 0.21, NMI 0.31) and Louvain (ARI 0.22, NMI 0.30), fail to distinguish the different cortical layers, leading to low clustering accuracies. These results underline the importance of effectively integrating spatial information into dimension reduction methods to optimize domain identification while preserving a moderate computational cost.

**Figure 2.**
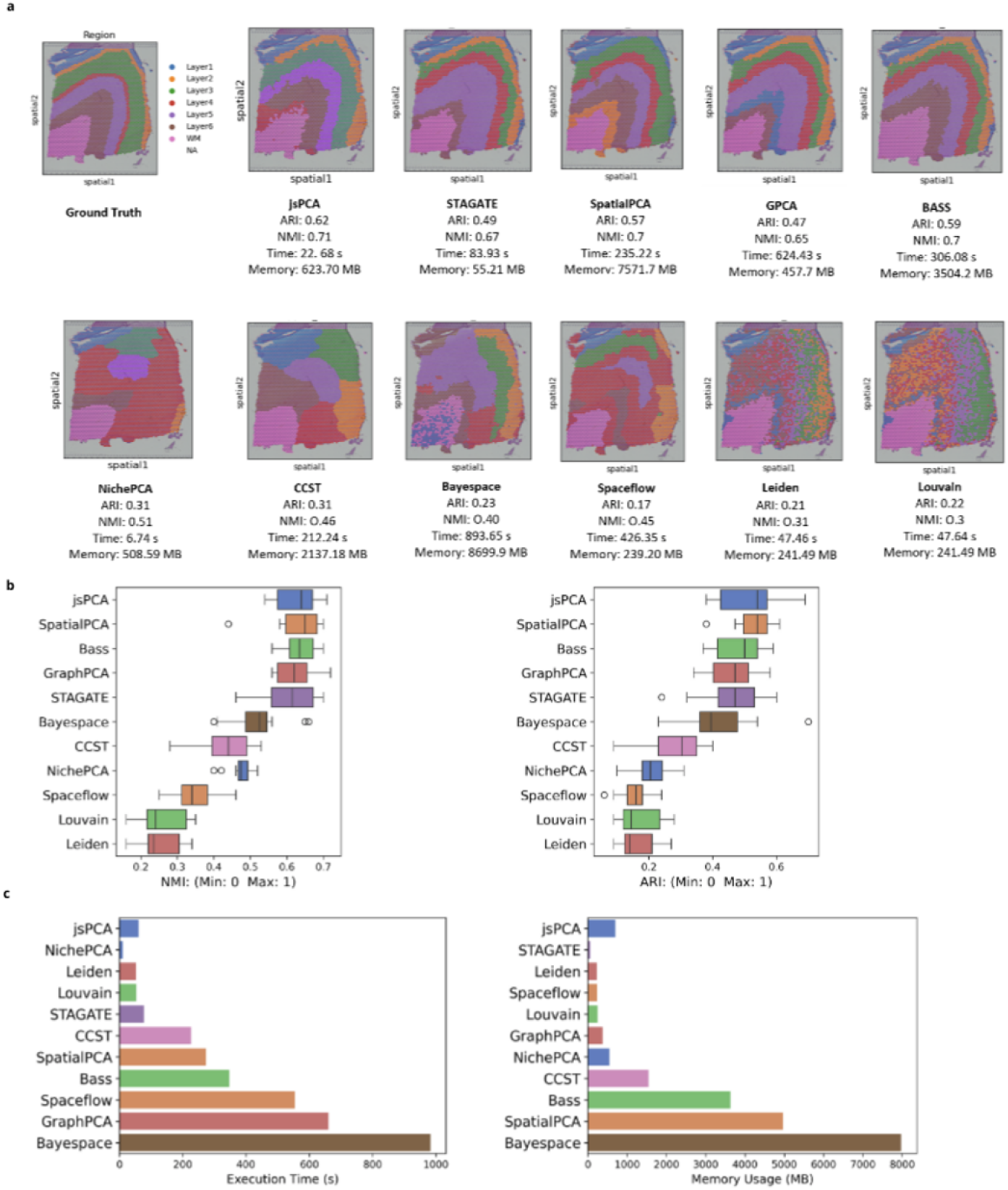
Performance of jsPCA for spatial domain identification in the human dorsolateral prefrontal cortex (DLPFC) 10x Visium dataset. **(a)** Spatial domain prediction results produced by different methods for slice 151673 of the DLPFC dataset, shown alongside the ground truth annotation of the six cortical layers and white matter (WM). ARI, NMI, runtime, and memory usage values are reported for each method. **(b)** Box plots showing clustering accuracy for all 12 DLPFC sections, measured by NMI (left) and ARI (right) across all methods. **(c)** Comparison of runtimes (left) and memory usage (right) across all methods.

**Figure 3.**
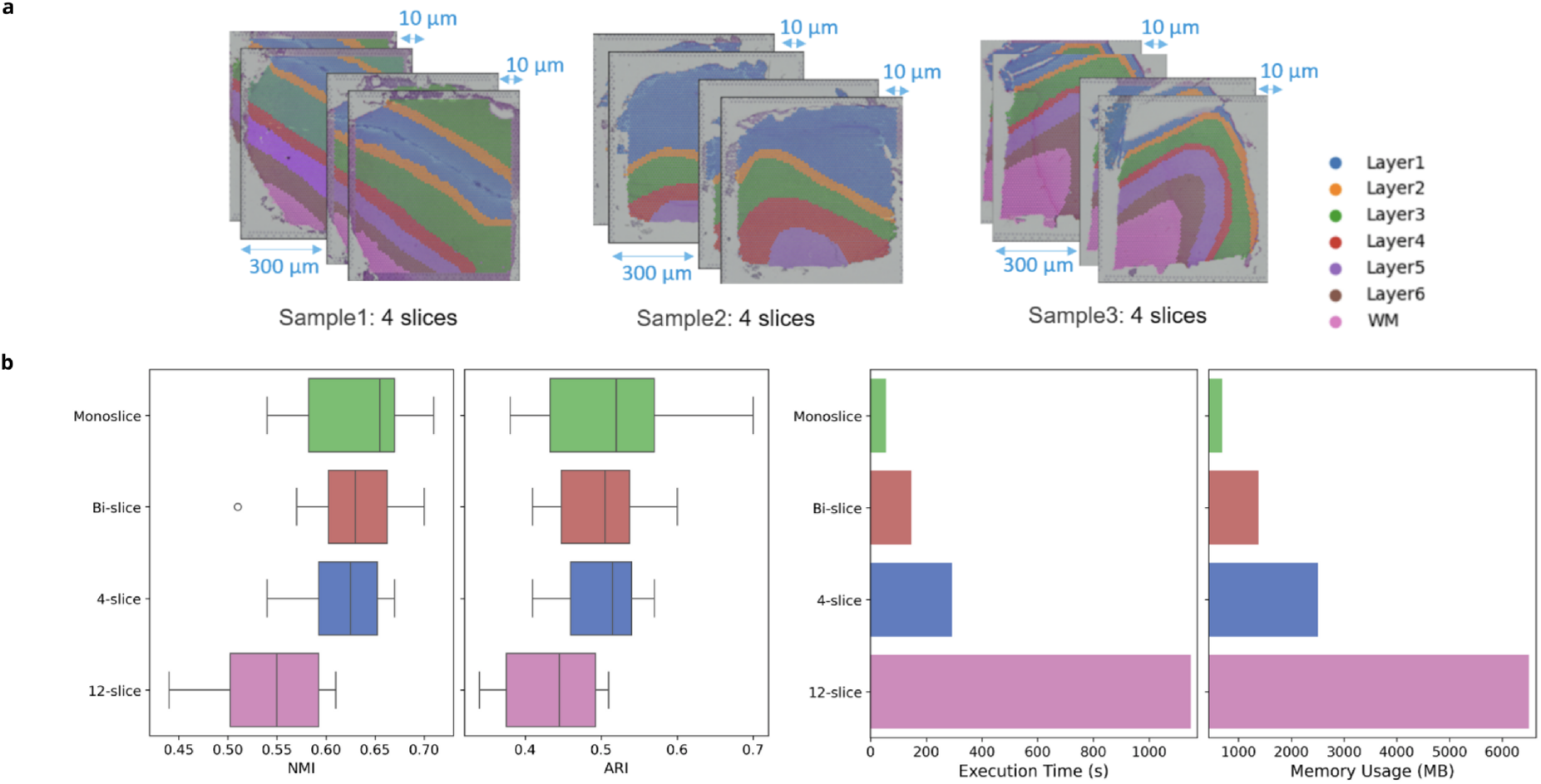
jsPCA performance across different configurations (monoslice, 2-slice, 4-slice and 12-slice) on the DLPFC dataset. **(a)** Distribution of the slices among three samples in the DLPFC dataset **(b)** Comparison of NMI and ARI scores and evaluation of runtime and memory usage.

In analyzing each of the sections contained in the entire database (Figure 2b), we noted that of all the spatial analysis methods compared in this study, jsPCA stands out for its outstanding balance between precision and reproducibility (remarkable stability and low variability of results). Its clustering performance, measured by ARI and NMI, reaches substantial median values, placing it in the top tier with SpatialPCA and Bass, but well above the other approaches. The real strength of jsPCA lies in its consistency, illustrated by the narrowness of its interquartile range for ARI and NMI. This low dispersion indicates that results remain reliable across different slices, unlike methods such as SpatialPCA (which has outliers) or STAGATE (which has higher variability). The total absence of outliers for jsPCA confirms its robustness in the face of special cases that could degrade the performance of other methods. Comparative analysis (Figure 2c) reveal that jsPCA also stands out for its computational efficiency, combining one of the shortest execution times (around 50 seconds) and moderate memory usage (800 MB), far below the requirements of SpatialPCA (5000 MB) or Bayespace (8000 MB).

After finding that, in its simplest setting, i.e. jsPCA on a single ST slice, our approach performs in a very fast and efficient way at accuracy level better or comparable to the top methods of the state of the art, we inquired the impact of considering multiple slices at the same time with jsPCA. Considering multiple slices at the same time has the advantage of predicting domains that are directly comparable between slices, as they are computed from a joint representation. Figure 3b shows the evolution of clustering performance, evaluated using the NMI and ARI indices, as a function of the number of slices. Accuracy is analyzed for different configurations: monoslice, 2-slice, 4-slice and 12-slice. While monoslice remains the most accurate, the 2-slice and 4-slice configurations still achieve good performance with only modest reductions. In contrast, the 12-slice configuration shows lower accuracy, which is expected since it merges slices coming from three different DLPFC samples, and these cross-sample differences reduce the consistency of the clustering. In terms of computational performances, however, there is a systematic increase of time and memory when increasing the number of slices considered together in jsPCA (Figure 3b).

jsPCA yields a representation of the data, the eigenvectors, that can be used independently of the dataset on which they have been computed. Therefore, jsPCA can in principle be used to predict a good embedding of an unseen dataset. To evaluate the prediction power of jsPCA, we performed a cross-slice analysis on the DLPFC dataset (Fig. 4a). For each sample, we trained jsPCA on three slices, learning the joint eigenvectors (principal components) and clustering parameters from the GMM model, and then used them to predict the domain labels of the remaining held-out slice. This process was repeated in a leave-one-out manner so that every slice served once as a test case. This approach makes it possible to test how well jsPCA generalizes from one slice to another within the same sample, showing whether it can handle structural differences while still producing consistent and reliable predictions. As shown in (Fig. 4b), the monoslice configuration reaches somewhat higher NMI score, but the cross-slice setting delivers strong and competitive results and even improve the ARI score. This indicates that jsPCA is able to generalize effectively across slices, handling structural variability without a major loss in accuracy.

**Figure 4.**
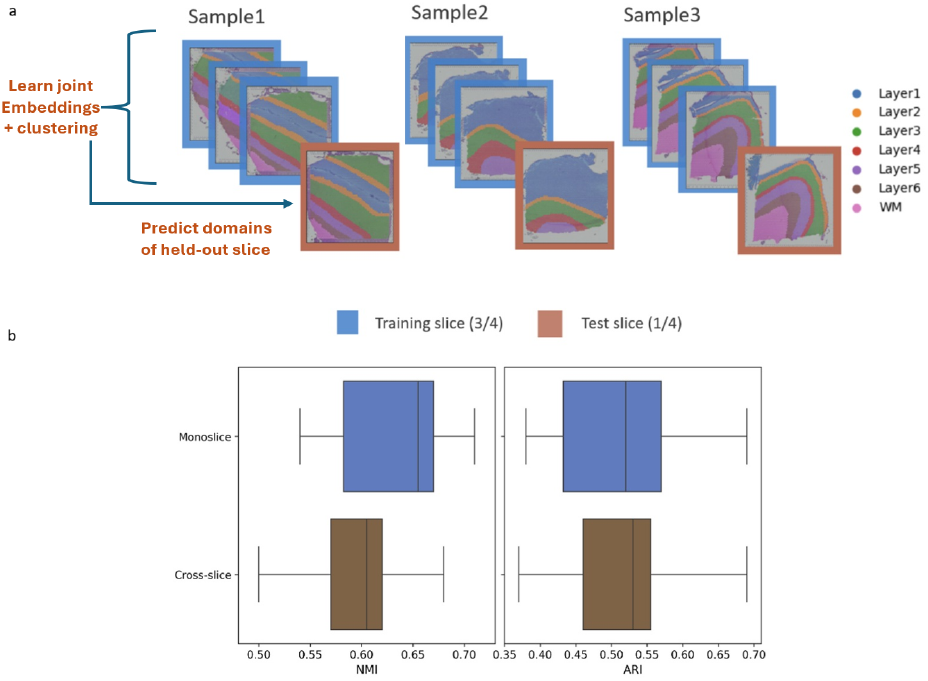
Cross-slice evaluation of jsPCA on the DLPFC dataset. **(a)** Schematic illustration of the cross-slice analysis. Each sample contains four slices. jsPCA was trained on three slices (blue boxes) to learn joint embeddings and clustering, and then used to predict the domains of the held-out slice (brown box). This procedure was repeated in a leave-one-out manner so that each slice served once as test data. **(b)** Quantitative comparison of monoslice (blue) and cross-slice (brown) configurations, measured by Normalized Mutual Information (NMI) and Adjusted Rand Index (ARI).

In Figure 5a, we compare the performance of jsPCA with several reference methods (STAGATE, SpatialPCA, GraphPCA, and BASS) for identifying spatial domains in the MOSTA dataset (E11.5 ES1). The domains obtained by jsPCA show a high degree of agreement with the manual annotation and achieves the best quantitative scores, with an NMI of 0.67 and an ARI of 0.48, outperforming the other methods. In Figure 5b, we present a monoslice analysis covering slices from developmental stage E9.5 through the first slice of stage E12.5. This subset was chosen because SpatialPCA and GraphPCA could not process the remaining slices of the dataset due to excessive memory requirements. Quantitative comparisons confirm the robustness of jsPCA, which achieves a strong balance between clustering accuracy and stability. The evaluation of runtime and memory usage demonstrates that jsPCA is computationally efficient relative to alternative methods, requiring on average only around 97 seconds and around 8.1 GB of memory, compared to several thousand seconds and much higher memory usage for SpatialPCA, GraphPCA, and BASS. In Figure 5c, we extend the monoslice analysis to include all slices of the dataset. jsPCA maintains strong clustering performance in terms of NMI and ARI, with stable results across the full range of slices. When considering runtime, jsPCA continues to be highly efficient, completing analyses much faster than STAGATE or BASS. In terms of memory, the requirements increase compared to the subset analysis, which is expected since the remaining slices contain a substantially larger number of spots. This higher memory demand explains why SpatialPCA and GraphPCA could not process these slices, whereas jsPCA successfully scaled to the entire dataset while preserving accuracy and robustness.

**Figure 5.**
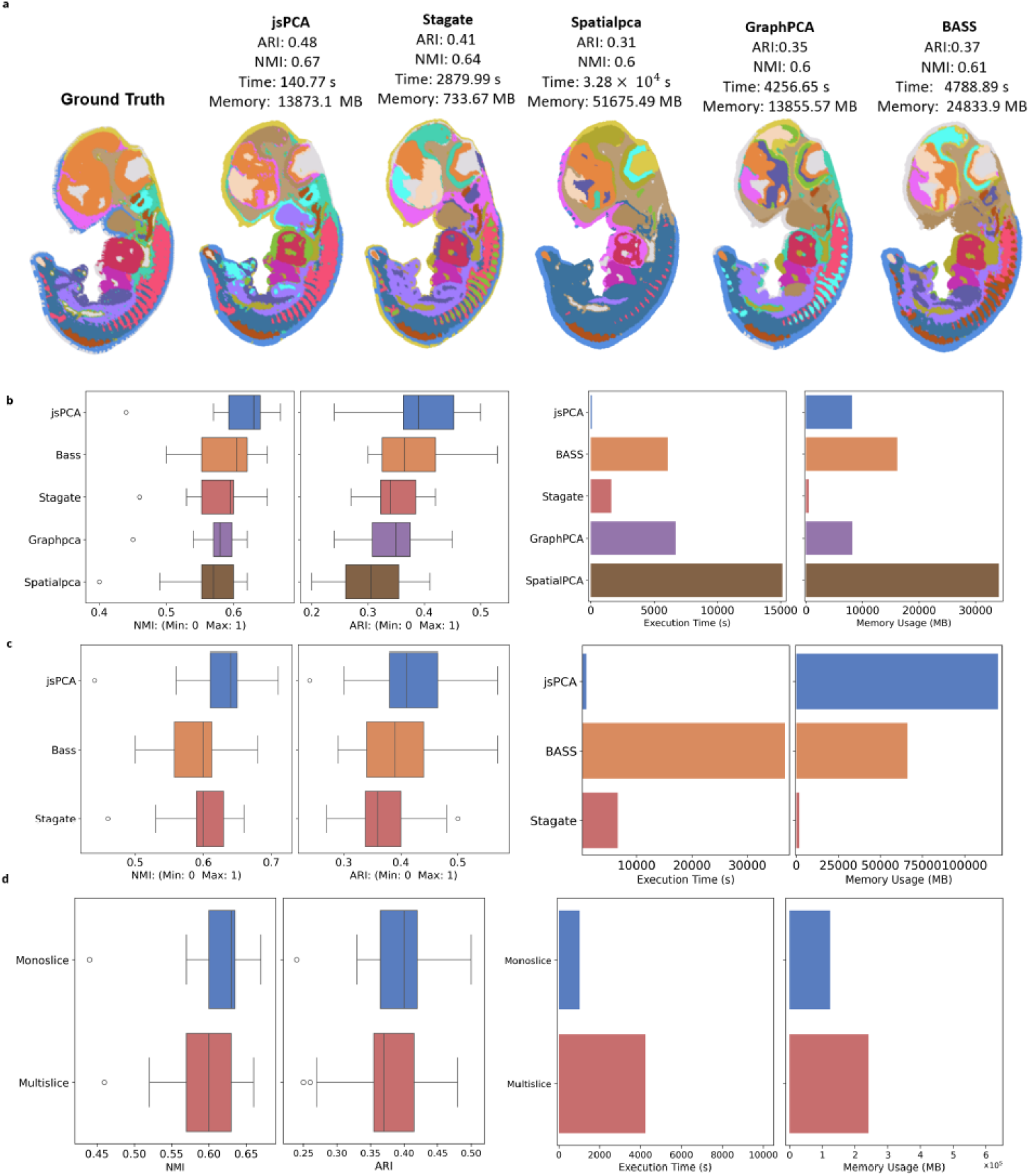
Performance of jsPCA for spatial domain identification in the mouse organogenesis spatio-temporal transcriptomic atlas (MOSTA) Stereo-seq dataset. **(a)** Ground truth obtained from [39]. Spatial domain identification results from the jsPCA, STAGATE, SpatialPCA, GraphPCA, and BASS methods on the E11.5 E1S1 sagittal section, shown alongside the manual reference annotation of major organs and tissues. ARI, NMI, runtime, and memory usage are reported for each method. **(b)** Comparative analysis on the first 14 sagittal sections of MOSTA, with box plots showing NMI (left) and ARI (right) scores for jsPCA, BASS, STAGATE, GraphPCA, and SpatialPCA, along with their runtimes and memory usage. **(c)** Additional evaluation on all sagittal sections of MOSTA for jsPCA, BASS, and STAGATE, using the same accuracy, runtime, and memory usage metrics. **(d)** Comparison of ARI and NMI scores, as well as runtime and memory usage, between monolayer and multilayer configurations across selected MOSTA slices (5 at E9.5, 4 at E10.5, 4 at E11.5, 6 at E12.5, and 4 at E13.5).

Figure 5d compares the performance of jsPCA on the MOSTA dataset in monoslice and multi-slice configurations. The monoslice configuration has overall slightly higher NMI and ARI scores, while the multi-slice configuration remains competitive, showing how robust jsPCA is in both cases. It also shows that analyzing multiple slices together requires more time and memory, which is predictable, but this increase in computational cost does not come at the expense of clustering quality. These results highlight jsPCA’s ability to deliver reliable and stable performance across different analysis settings.

Finally, to evaluate the ability of jsPCA to identify spatially variable genes (SVGs), we compared its outputs with those obtained from SPARK-X. The results show that jsPCA captures a substantial proportion of the SVGs identified by SPARK-X, particularly in its first component (denoted jsPCA1). In the DLPFC dataset (Fig 6a), jsPCA1 recovers about 82% of the SVGs detected by SPARK-X, while in the MOSTA dataset (Fig 6b) the overlap is nearly 79%. The distribution of jsPCA coefficients shows that the top-ranked genes contribute most of the spatial signal, which justifies selecting the top 3000 SVGs. When considering the top 3000 SVGs ranked by jsPCA1, the vast majority of SPARK-X genes are consistently present. These results highlight that jsPCA not only provides a simpler and more computationally efficient alternative to SPARK-X but also achieves a high level of agreement in the genes it identifies, supporting its reliability for large-scale spatial transcriptomics analyses.

**Figure 6.**
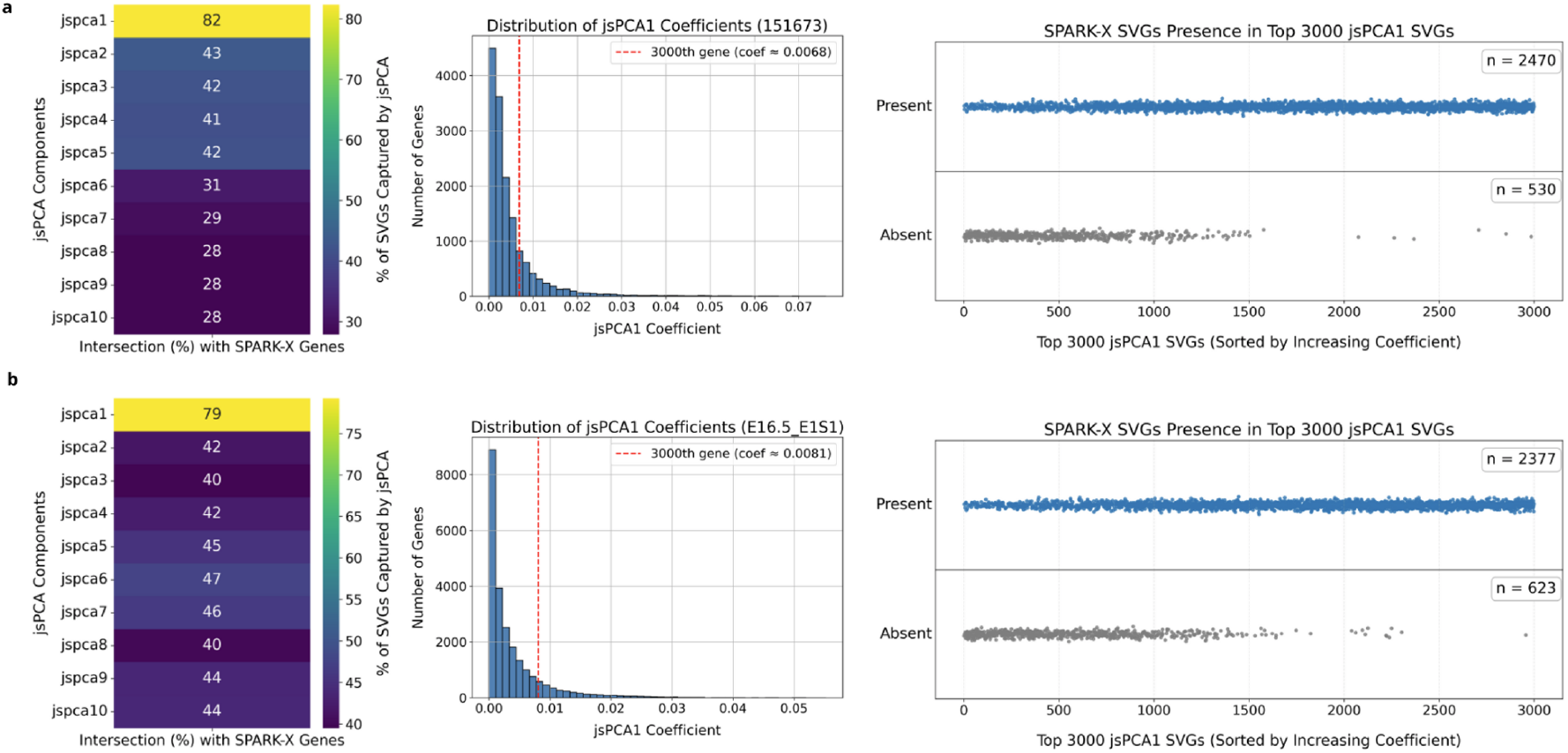
Detection of spatially variable genes by jsPCA: comparison with SPARK-X on **(a)** DLPFC (slice 151673) and **(b)** MOSTA (E16.15 E1S1). In the first column, we report the percentage of genes common between the 3000 most spatially variable genes identified by SPARK-X and the 3000 genes with the highest coefficients in each of top 10 eigenvectors. In the second column, we show the distribution of the coefficient in the top jsPCA eigenvector, the red dashed line represents the 3000th top coefficient. In the third column, we show how the genes common between SPARK-X and jsPCA are distributed in the jsPCA eigenvector, the x-axis represents the rank of the gene according to its coefficient value, the plots indicate if the gene is present or not in the SVGs identified by SPARK-X.

## Discussion

We introduced jsPCA, a computational framework for efficient and interpretable analysis of spatial transcriptomics data. By integrating gene expression covariance with spatial autocorrelation, jsPCA identifies both spatial domains and variable genes in a single formulation without the need for manual tuning of the spatial-expression trade-off. Compared with existing approaches, jsPCA consistently provides a favorable trade-off between accuracy, interpretability, and computational efficiency.

A key contribution of jsPCA is its ability to jointly analyze multiple tissue slices or samples without explicit spatial alignment. This contrasts with existing PCA-inspired methods, which typically operate on single slices or rely on distance kernels that do not generalize well across samples. We show that jsPCA generalizes across slices within the same tissue by performing accurate domain prediction on unseen slices. Moreover, jsPCA can be applied jointly to multiple samples, opening the way to integrative analyses of multi-sample ST datasets.

Our results also demonstrate that jsPCA is scalable to atlas-level datasets. On the Stereo-seq MOSTA dataset, jsPCA handled dozens of slices with tens of thousands of spots, whereas several leading methods failed due to excessive memory requirements. With the rapid emergence of ultra-high-resolution ST platforms such as Visium HD and whole-organ Stereo-seq, scalability is becoming a practical bottleneck; jsPCA offers a solution by maintaining efficiency without compromising interpretability.

Beyond benchmarking, jsPCA provides a framework for biological discovery. Because the principal components directly highlight spatially variable genes, jsPCA yields interpretable axes of variation that can be linked to known or novel tissue structures. This underscores the potential of jsPCA to uncover new biology in atlas-scale datasets.

Nevertheless, some limitations remain. jsPCA does not explicitly correct for batch effects, which can confound multi-sample comparisons. However, the modular nature of the framework allows for integration with established batch correction tools upstream of analysis [40]. Another limitation is that while jsPCA naturally extends to any spot- or cell-based dataset [26], its applicability to truly multi-modal omics data remains to be established. Future work will aim to extend jsPCA to joint spatial analysis across transcriptomic, proteomic, and metabolomic layers [41].

Overall, jsPCA provides an accessible, scalable, and biologically interpretable framework for spatial domain and gene identification in spatial transcriptomics. By enabling multi-sample joint analysis and scaling to datasets that currently exceed the reach of competing methods, jsPCA is positioned to become a standard tool for large-scale spatial omics studies, particularly in developmental biology and tissue atlas projects.

## Availability and Implementation

An open-source implementation of the jsPCA algorithm can be found on https://github.com/VILLOUTREIXLab/jsPCA.

## Acknowledgments

We thank Solène Song, Denis Puthier, Anaïs Baudot for fruitful discussions. This research was supported by Agence Nationale de la Recherche (ANR) project ANR-CPJ12022-VILLOUTREIX-U125. It benefited from access to the Core Cluster of the Institut FranÇais de Bioinformatique (IFB).

## Supplementary Text 1 - Computation of jsPCA for large matrices

To compute jsPCA eigenvectors for a collection of ST datasets, i.e. a collection of expression matrices and normalized adjacency matrices {*X*_*j*_, *L*_*j*_}, we need to perform a joint diagonalization of the corresponding spatial covariance matrices. This can be summarized by finding the common set of eigenvectors *P* such that

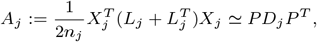

The matrices *P* and {*D*_*j*_} can be obtained by solving

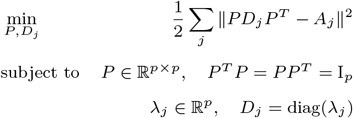

Given the large size of the matrices handled in ST, the joint diagonalization of the spatial covariances matrices {*A*_*j*_} cannot be obtained by classical approaches such as Jacobi-like techniques because they require too much memory [42]. Instead, we implemented the following approach.

### Independent truncated diagonalizations

First, the truncated diagonalizations of order *K* of each *A*_*j*_ are computed as 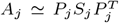. In our experiments, we use values of *K* ranging from 10 to 50, hence *K* ≪ *p*. For *K* ≪ *p* the truncated diagonalization can be efficiently computed using an Implicitly Restarted Lanczos Method which is implemented in the scipy library [43]. The key feature of this method is its *matrix-free* operation meaning that it only uses matrix-vector products with *A*_*j*_ and does not require a matrix built in memory. However, since *p* is large and the matrices *A*_*j*_ are dense, directly computing the eigensystem from *A*_*j*_ is particularly inefficient (considering the building time and the dense matrix-vector products). Fortunately the matrices *X*_*j*_ and *L*_*j*_ are very sparse. Implementing an efficient matrix-vector product with *A*_*j*_ can therefore be performed by successive sparse matrix-vector products, that is implementing 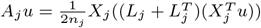 for *u* ∈ ℝ^*p*^

### Joint approximate diagonalization

Given the eigensystems {(*P*_*j*_, *D*_*j*_)}_*j*_, the joint diagonalization can now be modeled as

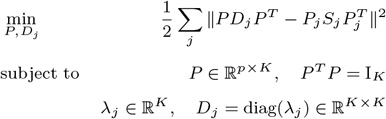

With an argument similar to [44] this problem can be shown to be equivalent to

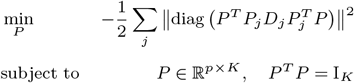

which can be solved by a gradient descent on the Stiefel manifold [27, 28] (modeling the orthogonality constraint *P* ^*T*^ *P* = I_*K*_). Following again [44], an initialization close to a minimizer can be obtained by computing the *K* largest left singular vectors of the matrix (*P*_1_*S*_1_, …, *P*_*j*_ *S*_*j*_, …) (i.e. concatenating the columns of the {*P*_*j*_ *S*_*j*_}) and by letting 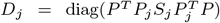. In our experiments, we performed 100 steps for the gradient descent.

To further justify this choice, we performed an additional convergence analysis of the joint diagonalization step (Figure S9). Specifically, we tracked the evolution of the cost *J* ^(*t*)^, which is equal to the quantity 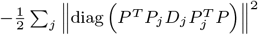 evaluated at iteration *t* of the gradient descent and the normalized convergence gap (*J* ^(*t*)^ − *J* ^*⋆*^)*/*(*J* ^(0)^ − *J* ^*⋆*^) across iterations, for both random and SVD initializations. The experiment was performed on two slices from the DLPFC dataset (151673 and 151674), comprising 3639 and3673 spots respectively, with 15026 genes per slice. The results show that the cost decreases rapidly during the first few iterations and then reaches a stable plateau. In particular, most of this decrease occurs during the first 100 iterations, after which the improvement becomes negligible. While both initialization methods reach similar final values, the initial point provided by the SVD is very close to the optimum, roughly saving 4000 iterations compared to a random initialization. This is substantiated by the normalized convergence curve showing that the steepest descent only marginally improves the cost of the current iterate. For this reason, we use SVD initialization in all experiments together with 100 gradient descent iterations.

## Supplementary Text 2 - Formal comparison between jsPCA and spatial-PCA like approaches

We review in this section the differences between various formulations of *spatial PCA* applied to spatial transcriptomics that have been proposed recently, with harmonized notations.

We consider a matrix of gene expression measurements for *p* genes in *n* spatial locations of a tissue, denoted by *X* ∈ ℝ^*n*×*p*^. These measurements have spatial coordinates that are denoted by *S* = {*s*_*i*_ ∈ ℝ^*k*^, *i* ∈ {1, .., *n*}} where *k* = 2 or 3.

Recall the variational formulation of PCA, which consists in finding a *d*-dimensional projection *Z* ∈ ℝ^*n*×*d*^ such that:

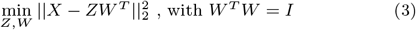

This optimization problem can be solved analytically. The analytical solution leads to *Z* = *XW* where *W* represents the top-*d* eigenvectors of the empirical covariance matrix *X*^*T*^ *X*.

PCA however does not take into account the spatial information *S*. Several methods have been proposed to incorporate spatial information in the context of spatial transcriptomics data analysis. We will distinguish between two families of approaches, the approaches based on the distance between spots and the approaches based on a spatial graph. In that case, the locations are turned into an undirected graph with the nearest neighbor algorithm, we denote by *C* the corresponding adjacency matrix. *C*_*i,j*_ = 1 if *j* is in the neighborhood of *i*, 0 otherwise.

### Distance-based spatial PCA *Spatially aware dimension reduction for spatial transcriptomics [15]*

The expression matrix *X* is assumed to have been normalized through variance stabilizing transformation and further scaled for each gene to have zero mean and unit standard deviation.

The goal is to infer a *n* × *d* factor matrix *Z* that represents a low dimensional embedding of *X*. The approach is based on the following latent factor model:

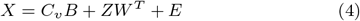

where *C*_*v*_ is an *n* × *q* matrix of covariates, and *B* is a *q* × *p* matrix of corresponding coefficients, here *q* is assumed to be equal to 1. *W* is a *p* × *d* factor loading matrix and *E* is an *n* × *p* matrix of residual errors, with elements *E*_*i,j*_ following independent normal distribution with mean zero and variance 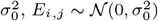.

*W* is assumed to have orthonormal columns such that *W* ^*T*^ *W* = *I*. For the *d* factors *Z*_*l*_, i.e. the vectors of dimension *n* composing the factor *Z* matrix, the assumption is that they follow a multivariate normal distribution:

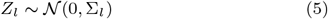

with Σ_*l*_, the *n* × *n* covariance matrix modeling the correlation among the spatial locations. Σ_*l*_ has for elements 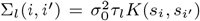 with Gaussian kernel 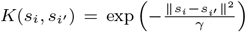. *τ*_*l*_ and *γ* are two hyperparameters controlling the scale of the factor value relative to the residual error and the strength of the spatial correlation, respectively. *γ* is set numerically, in relation to the range of values in the data. *τ*_*l*_ = *τ* is assumed constant across factors.

Maximum likelihood-based inference predicts the factor loading matrix *W* and the factor matrix *Z* together with the hyper-parameters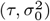.

The spatial domains are obtained by computing a clustering algorithm on *Z*, in this case Walktrap algorithm and Louvain algorithm for clustering analysis.

### Randomized Spatial PCA (RASP): a computationally efficient method for dimensionality reduction of high-resolution spatial transcriptomics data [32]

This approach first computes a randomized PCA on the expression matrix *X* with rank *d* and computes *W*, the matrix containing the top-*d* principal components of the covariance. The data is projected on the principal components and stored in a matrix *Z* = *XW*. In a second step, the spatial information is taken into account with the connectivity matrix *C* of the nearest neighbors graph. From this connectivity matrix, a distance matrix *D* is computed with diagonal elements set to a value *α*. The matrix is inverted with a power *β*, and stored in a matrix *D*_inv_ = (*D*^−1^)^*β*^ which is subsequently normalized column-wise. The final representation is obtained by computing the product *Z*_*s*_ = *D*_inv_*Z* = (*D*^−1^)^*β*^*XW*, which gives a spatially smoothed version of PCA-derived projection.

### Graph-based spatial PCA *GraphPCA: a fast and interpretable dimension reduction algorithm for spatial transcriptomics data [16]*

The expression matrix *X* is assumed to have been normalized by analytic Pearson residuals proposed by [45] and further scaled for each gene to have zero mean and unit standard deviation. Then, the top 3000 spatial variable genes of the SPARK package are selected.

GraphPCA takes physical distance into account by including a graph regularization to the variational PCA (3) leading to

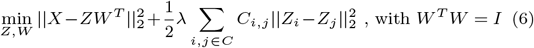

where *λ* is a hyperparameter controlling the reconstruction error versus the smoothness of the projections on the underlying spatial neighborhood graph. This optimization problem can be rewritten as

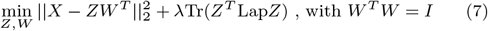

where Lap = *D* − *C* is the Laplacian matrix of the spatial neighborhood graph, with *D* the diagonal matrix containing the degree of the nodes in the graph *C*.

The solution of equation (7) obtained with Lagrange multiplier is *Z*^*^ = (*I* + *λ*Lap)^−1^*XW* and from this it follows that *W* ^*^ is the collection of top-*d* eigenvectors of the graph-revised covariance matrix *X*^*T*^ (*I* + *λ*Lap)^−1^*X* = *X*^*T*^ (*I* + *λ*(*D* − *C*))^−1^*X*.

### PCA-based spatial domain identification with state-of-the-art performance [33]

The preprocessing step for the expression data consists in normalization and log-transform per cell. In the same way as in jsPCA and graphPCA, for nichePCA [33], the spatial information is encoded in a *k*-nearest neighbor algorithm. The joint spatial and expression representation is obtained by computing the mean gene expression of a cell and its neighbors. Classical PCA is computed on this joint representation to obtained a spatially aware low-dimensional representation.

### Comparison with jsPCA

We have reviewed here recent approaches aimed at adapting classical PCA to spatially aware dimensionality reduction for spatial transcriptomics data. While they revolve around similar ideas, the mathematical formulations differ, leading to different numerical solutions.

jsPCA is based, for a single ST sample, on the diagonalization of the following spatial covariance which is obtained as the product of the spatial autocorrelation and the gene expression covariance:

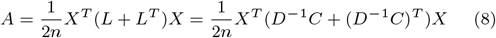

The corresponding embedding is obtained as *Z* = *XW*, where *W* correspond to the top-*d* eigenvectors of the spatial covariance A.

From this analytical solution, we can directly see that this formulation is quite different from the one called spatialPCA in [15], equations (4) and (5). Compared to jsPCA, spatialPCA is distance-based and relies on a Gaussian kernel for the spatial covariance. Moreover, the factors of spatialPCA [15] can be found with maximum-likelihood based inference. Their complete derivation is quite heavy and can be found in [15] supplementary material. It is noticeable that the influence of the spatial information is controlled by the hyperparameter *τ*.

Similarly for the method RASP [32], the optimal representation is obtained by *Z*_*s*_ = (*D*^−1^)^*β*^*XW*, where *D* is a spatial distance matrix, and *W* corresponds to the eigenvectors of the PCA. Analytically, the solution is quite different from jsPCA and rely on a parameter *β* to control the influence of space, as well as several heuristic steps.

GraphPCA is closer to jsPCA as it is a graph-based approach, relying on a Laplacian regularization of the variational formulation of PCA [46]. The corresponding graph-revised covariance matrix reads *X*^*T*^ (*I* + *λ*(*D* − *C*))^−1^*X*, to be compared with the spacial covariance of jsPCA in equation (8). With this rewriting of the two covariance matrices, it is clear that the two approaches have rather different grounds.

Finally, NichePCA computes the mean of the expression of spatial location and its direct neighbor as the new feature on which to apply PCA. Although it exploits the direct neighbors information, this approach does not capture the full extent of cross terms that jsPCA captures by computing the spatial autocorrelation on the nearest neighbors graph.

### Statistical significance analysis of jsPCA performance

To assess whether the observed differences could be explained by chance, we performed statistical comparisons between jsPCA and the other methods across DLPFC and MOSTA datasets using both parametric (paired t-test) and non-parametric (Wilcoxon signed-rank) tests, based on ARI and NMI scores. The detailed results are provided in Tables S2-S7 and Figures S1-S3.

The results show that jsPCA consistently outperforms several methods, including Leiden, Louvain, NichePCA, and Spaceflow, with statistically significant differences and large effect sizes for both ARI and NMI. These improvements are stable across datasets and are confirmed by both statistical tests. When compared to stronger baselines such as BASS, STAGATE, GraphPCA, and SpatialPCA, jsPCA achieves overall competitive performance, and in several cases outperforms them. In particular, jsPCA shows significant gains on the MOSTA dataset, while the differences are smaller and sometimes not statistically significant on the DLPFC dataset. This indicates that jsPCA is able to match or exceed their performance depending on the dataset.

In summary, these results indicate that the improvements observed with jsPCA are not due to random variation, but reflect a robust and competitive performance across different datasets and evaluation metrics.

### Biological interpretation of jsPCA principal components

To interpret the signals captured by the principal components from a biological perspective, we examined how known marker genes, as reported in the DLPFC study [37] are represented within each component (Figure S4 and Table S8). The spatial expression patterns of these marker genes in the ground truth data are shown in Figures S5-S6 and are used as an external reference for biological interpretation. The components capture distinct and meaningful spatial patterns. jsPC1 clearly reflects white matter (WM), as indicated by the strong contribution of MBP (rank 1) and MOBP (rank 6), while layer specific markers such as CUX2, RORB, and PCP4 are not enriched. jsPC2 highlights superficial cortical regions, which is supported by the enrichment of HPCAL1 (rank 2) and CUX2 (rank 38), consistent with layer 2 expression. In addition, SNAP25 (rank 9), a marker of gray matter (GM), further supports the cortical origin of this component. jsPC4 captures a mixed laminar signal, with contributions from both superficial and deep layers, as shown by HPCAL1 (rank 35, L2) and KRT17 (rank 24, L6). jsPC5 is primarily associated with layer 1 via AQP4 (rank 11), with a smaller additional contribution from deeper layers, as suggested by KRT17 (rank 27, L6). jsPC6 shows a weak but constant signal from layer 5, corroborated by PCP4 (rank 10) and a slight spatial enrichment in L5. In contrast, jsPC7 exhibits a strong, well-defined laminar structure dominated by the deeper layers (L4–L6), with high-ranking markers such as PCP4 (rank 1, L5), KRT17 (rank 5, L6), TRABD2A (rank 7, L1), and RORB (rank 12, L4), while CUX2 (rank 12411, L2) is not represented. The higher-order components, jsPC8 and jsPC9, are dominated by IGKC (rank 1 in jsPC8 and rank 4 in jsPC9) and reflect sparse, localized non-neuronal or immune-related signals. Finally, jsPC10 detects a weak mixed signal, with contributions from AQP4 (rank 26) and HBB (rank 27), suggesting a combination of non-neuronal expression or vascular origin, with a small contribution from layer 1. Overall, these results show that jsPCA decomposes the data into components that reflect both major anatomical structures, such as white matter and cortical layers, and secondary or non-neuronal sources of variation.

### Effect of Spatial Refinement and Threshold Sensitivity

We evaluated clustering performance both before and after applying the spatial refinement step, as well as across different refinement thresholds. As shown in Figure S8, the jsPCA embedding already gives strong NMI and ARI scores without any refinement, indicating that the main structure is well captured on its own. Applying refinement leads to a small but consistent improvement, mainly by smoothing local inconsistencies, but the overall gains remain limited. This suggests that most of the performance comes from the jsPCA embedding rather than this post-processing step. We also find that performance is robust to the choice of threshold. Moderate values (0.3–0.5) tend to give slightly higher median scores, while more stringent thresholds (≥ 0.7) do not provide further improvements. Overall, the variation across thresholds is small, indicating that the method does not depend on careful tuning of this parameter.

**Figure S1.**
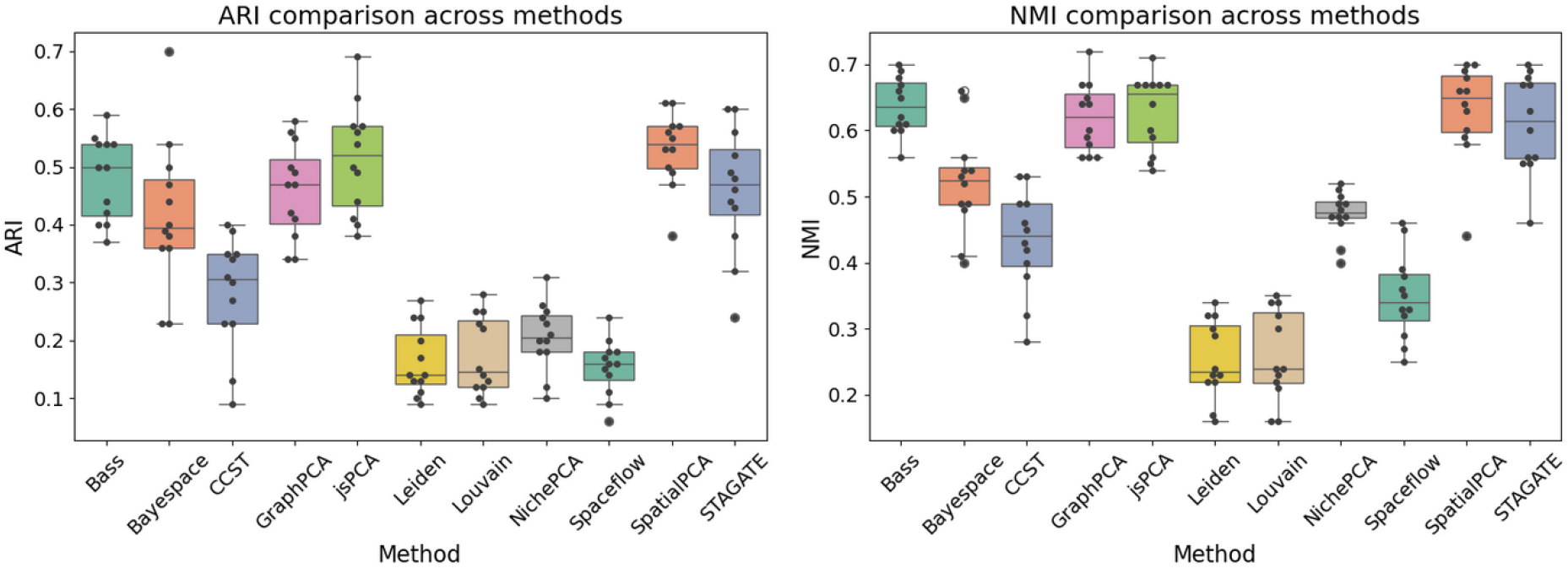
ARI and NMI distributions across methods on the DLPFC dataset. Statistical differences between jsPCA and other methods were assessed using both a paired t-test and a Wilcoxon signed-rank test (see Table X). Both tests lead to identical boxplot visualizations, so, a single figure is shown.

**Figure S2.**
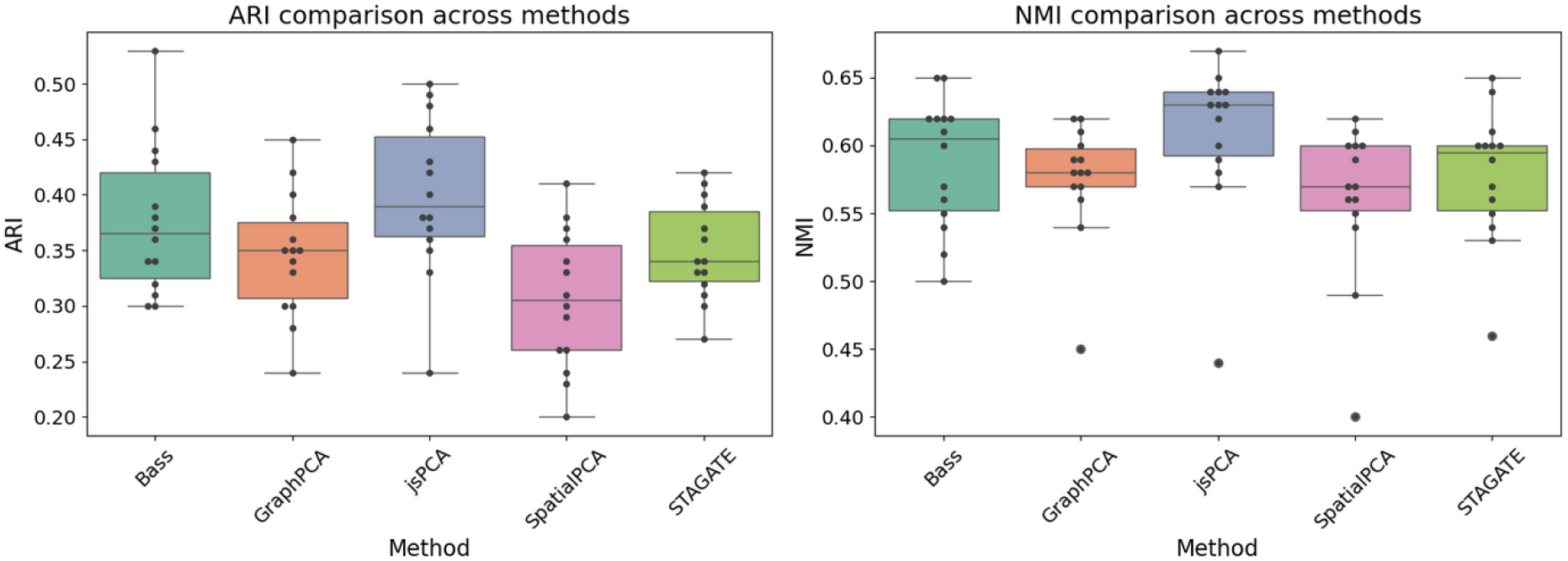
ARI and NMI distributions across benchmark methods on the first 14 sagittal sections of the MOSTA dataset. Statistical differences between jsPCA and other methods were assessed using both a paired t-test and a Wilcoxon signed-rank test (see Table X). Both tests produce identical boxplot visualizations, so, a single figure is shown.

**Figure S3.**
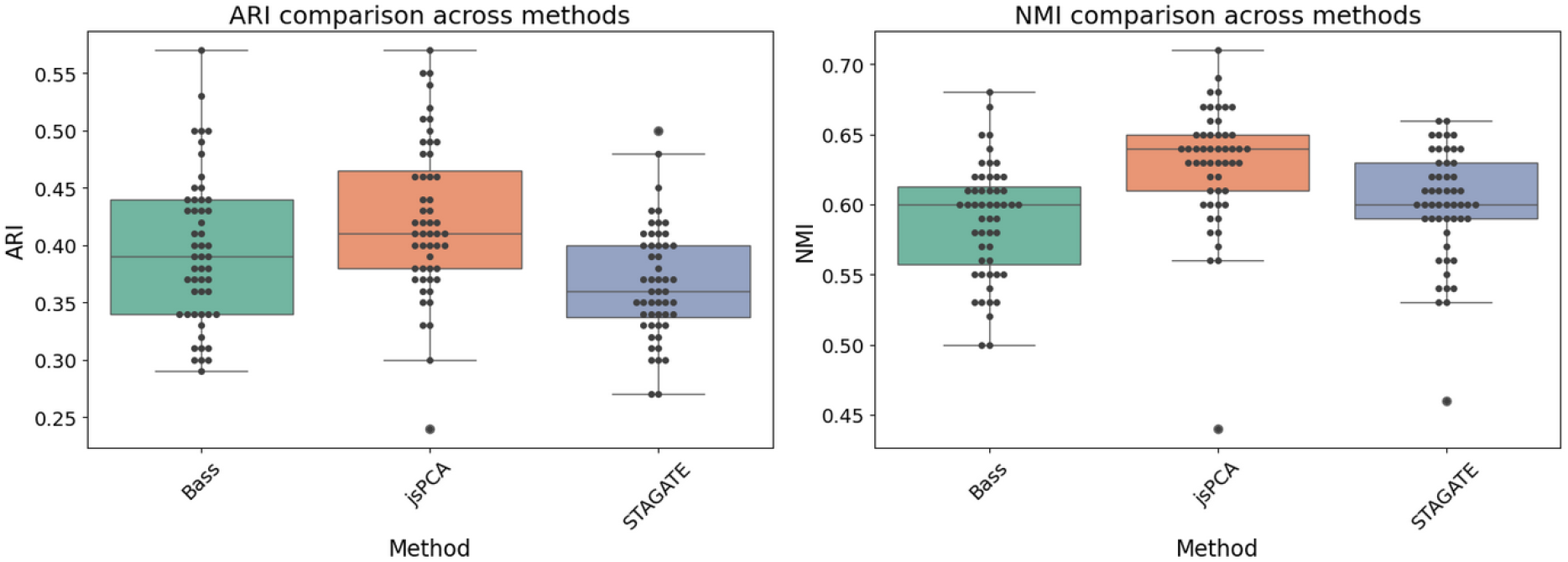
ARI and NMI distributions across jsPCA, BASS, and STAGATE on the whole MOSTA dataset. Statistical differences between jsPCA and the other methods were assessed using both a paired t-test and a Wilcoxon signed-rank test (see Table X). Both tests produce identical boxplot visualizations, so, a single figure is shown.

**Figure S4.**
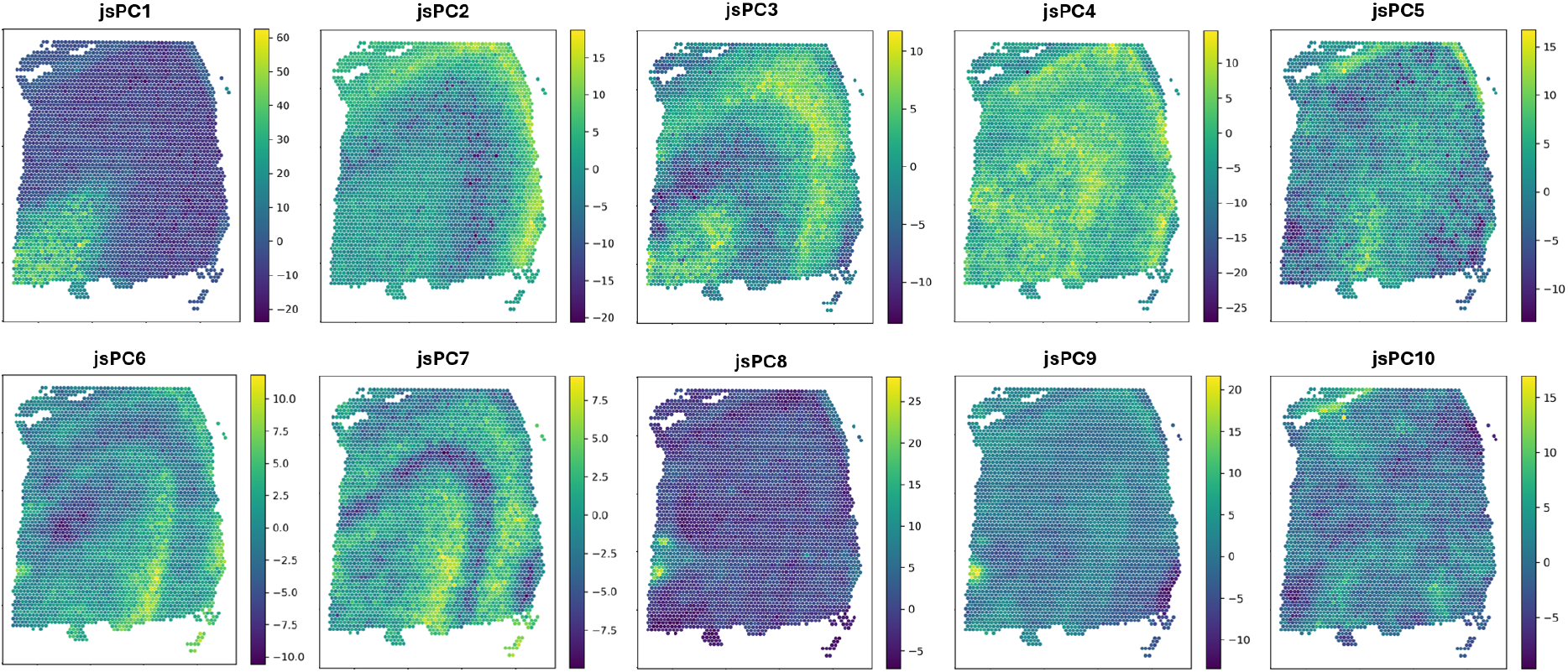
Visualization of each of the 10 principal components of jsPCA for the slice 151673 of the DLPFC dataset.

**Figure S5.**
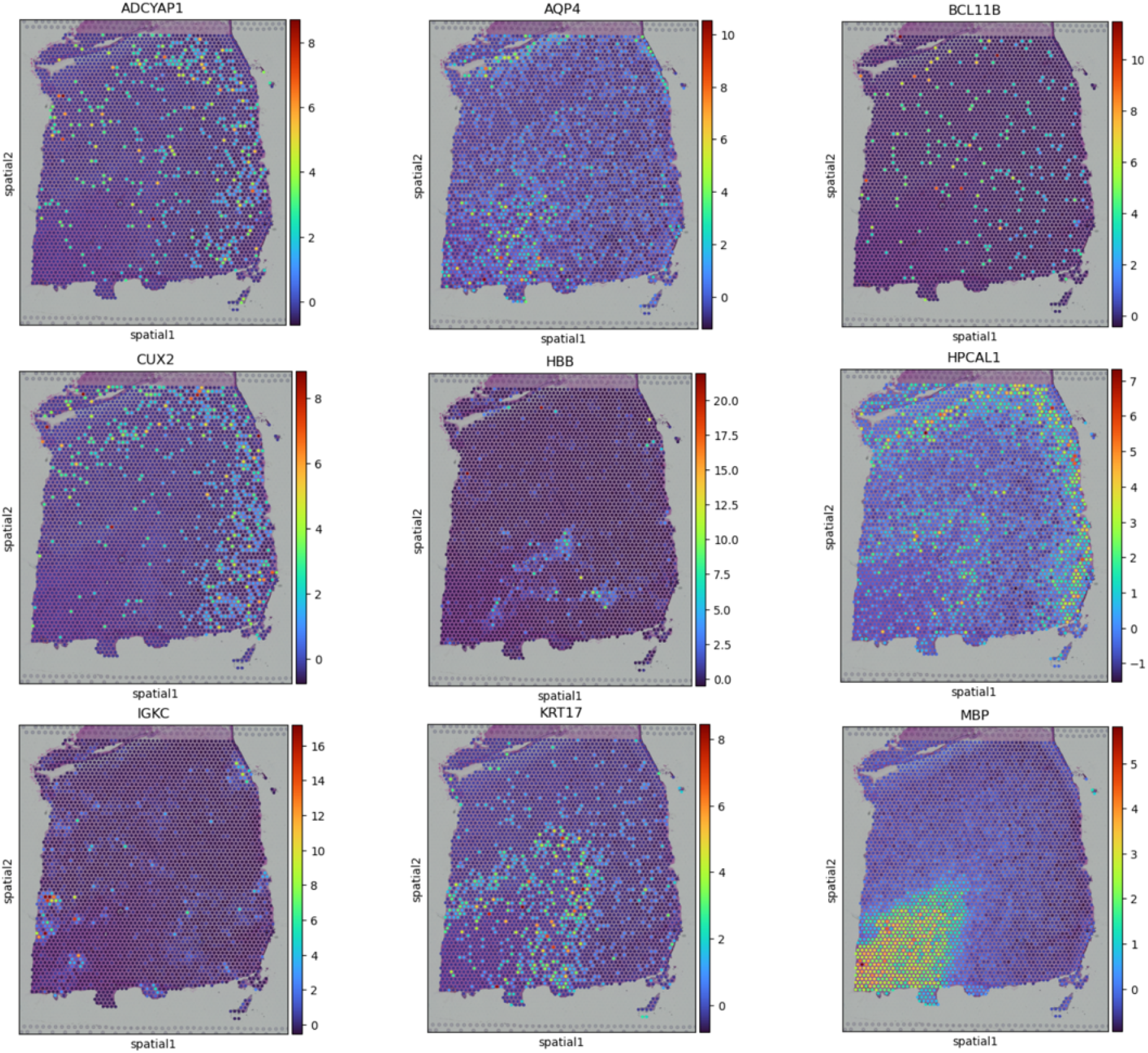
Spatial expression patterns of marker genes (ADCYAP1, AQP4, BCL11B, CUX2, HBB, HPCAL1, IGKC, KRT17, MBP) in the ground truth tissue for slice 151673 of the DLPFC dataset.

**Figure S6.**
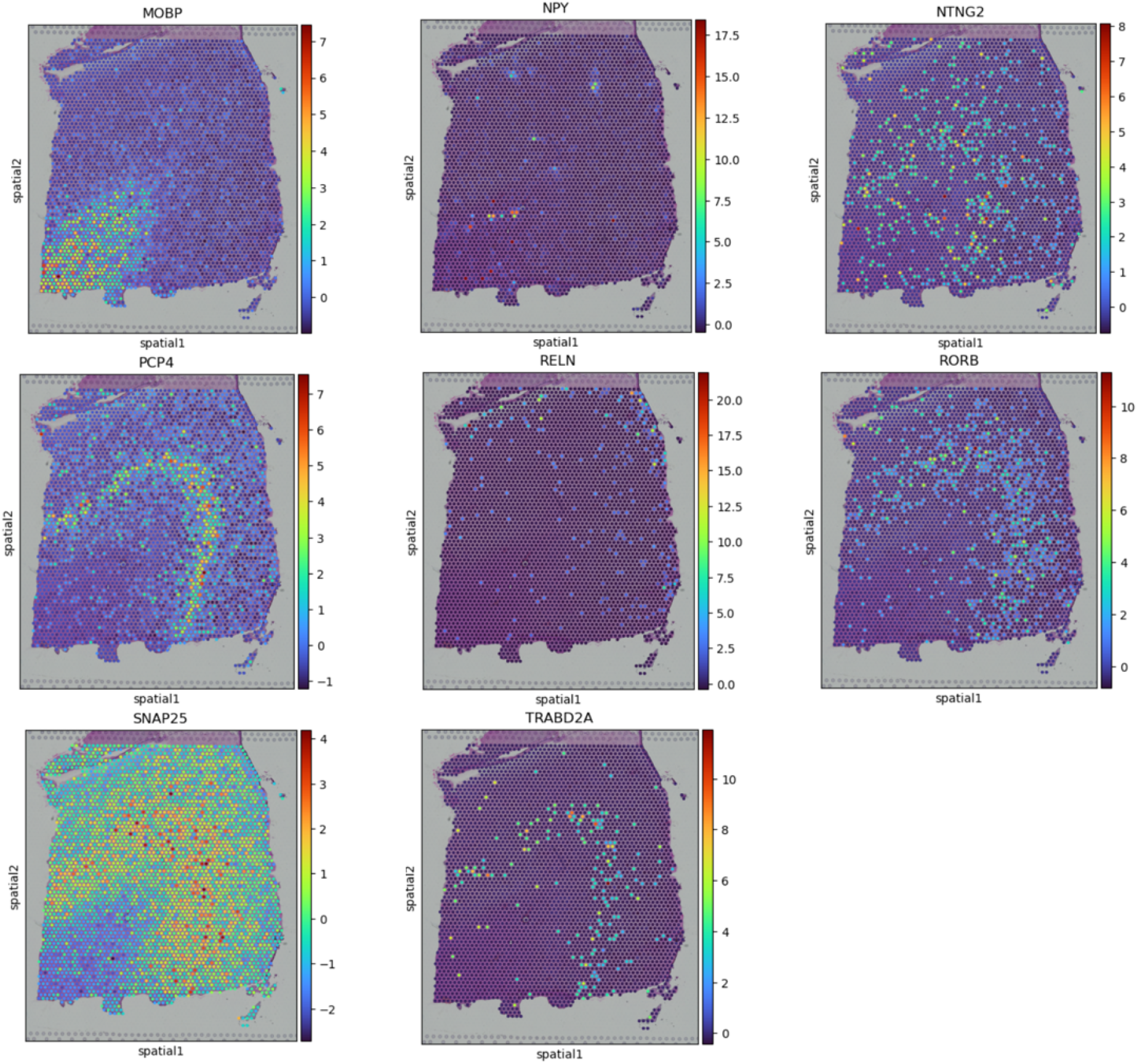
Spatial expression patterns of marker genes (MOBP, NPY, NTNG2, PCP4, RELN, RORB, SNAP25, TRABD2A) in the ground truth tissue for slice 151673 of the DLPFC dataset.

**Figure S7.**
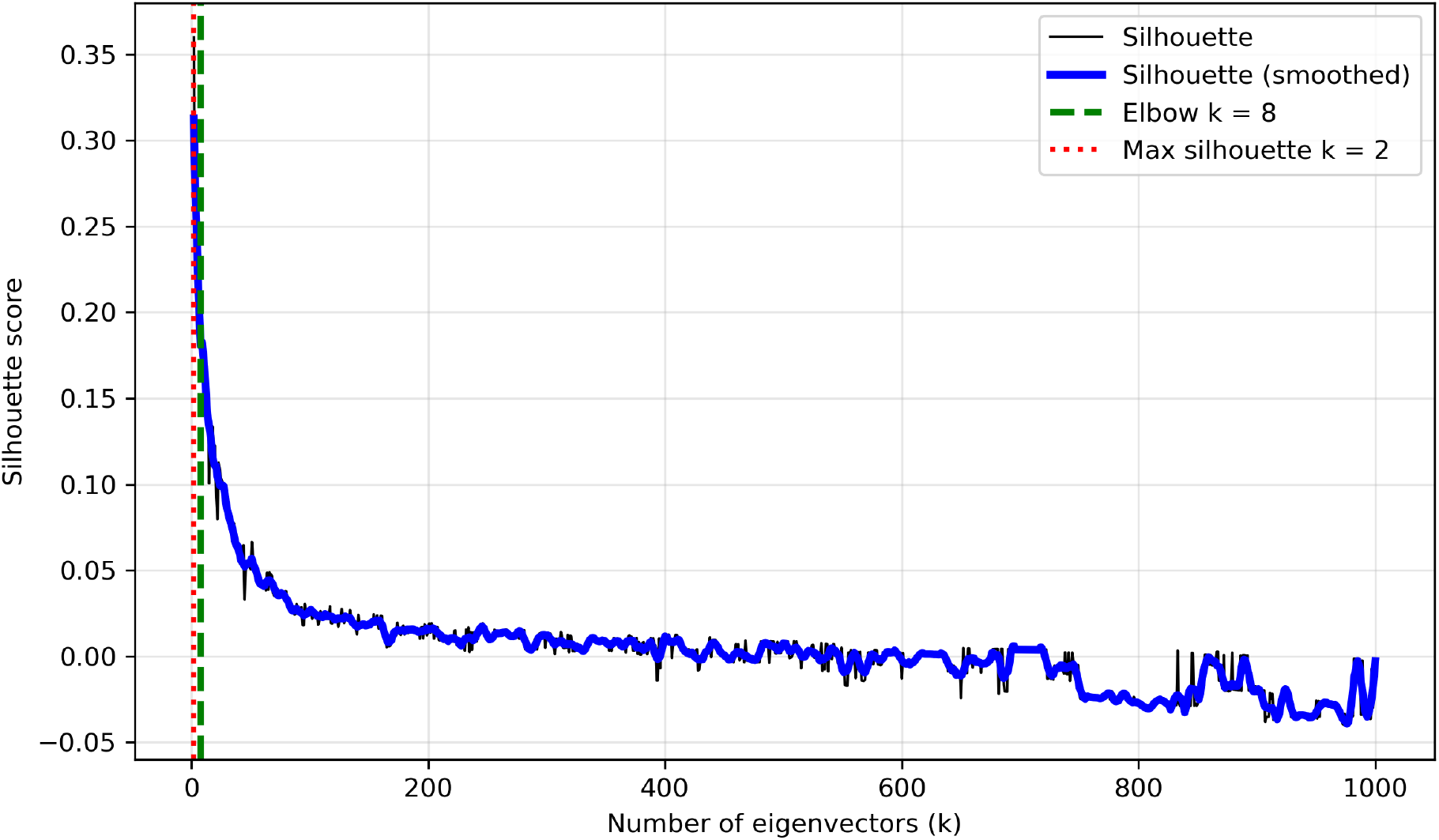
Visualization of the silhouette score as a function of the number of retained eigenvectors (k) for the slice 151673 of the DLPFC dataset. The raw (black) and smoothed (blue) curves are shown, along with the selected elbow point (k=8, green dashed line) and the maximum silhouette value (k=2, red dotted line).

**Figure S8.**
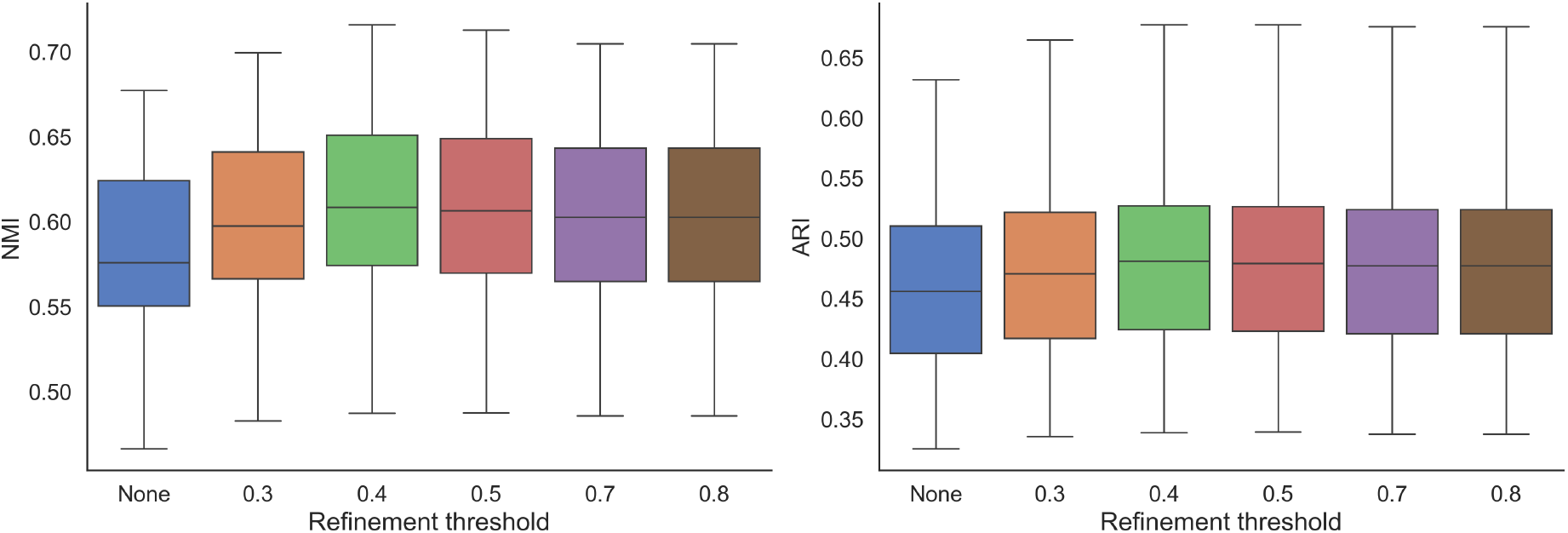
Clustering performance across refinement thresholds for the DLPFC dataset.

**Figure S9.**
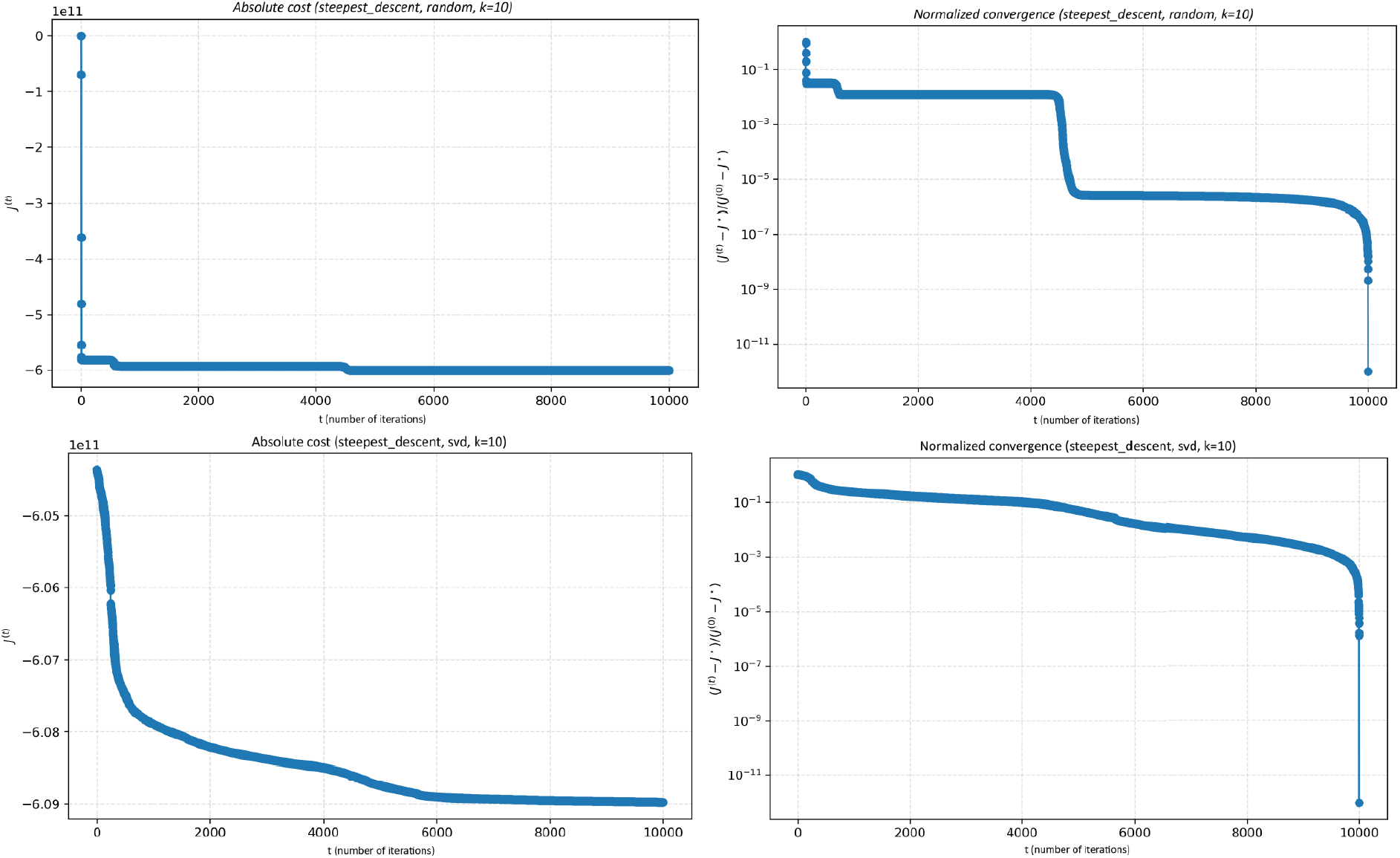
Convergence behavior of the steepest descent algorithm under different initialization strategies (random vs. SVD).

**Table S1.** Overview of gene selection procedures across methods.

| Method | Genes selected in benchmark |
| --- | --- |
| jsPCA | All genes passing basic prevalence filtering (genes expressed in < 20 cells) |
| STAGATE | Top 3000 HVGs |
| SpatialPCA | 3000 HVGs, 3000 SVGs, or all genes via an intermediate PCA step (3000 PCs) |
| GPCA | 3000 SVGs selected by SPARK |
| BASS | 3000 SVGs selected by SPARK-X |
| NichePCA | Removing all genes expressed in < 5 cells |
| CCST | All input genes (no feature selection step) |
| Bayespace | Top 2000 HVGs |
| Spaceflow | Top 3000 HVGs |
| Leiden | All genes passing basic prevalence filtering (genes expressed in < 20 cells) |
| Louvain | All genes passing basic prevalence filtering (genes expressed in < 20 cells) |

**Table S2.** Paired t-test comparison of jsPCA against other benchmark methods based on ARI and NMI, using the DLPFC dataset.

| Metric | Method | p-value | Mean diff |
| --- | --- | --- | --- |
| ARI | Bass | 0.0748 | 0.0317 |
| ARI | Bayespace | 0.0576 | 0.0975 |
| ARI | CCST | 5.55E-06 | 0.2317 |
| ARI | GraphPCA | 0.0853 | 0.0550 |
| ARI | Leiden | 1.52E-07 | 0.3508 |
| ARI | Louvain | 6.42E-07 | 0.3408 |
| ARI | NichePCA | 3.63E-08 | 0.3075 |
| ARI | STAGATE | 0.0400 | 0.0542 |
| ARI | Spaceflow | 5.83E-07 | 0.3608 |
| ARI | SpatialPCA | 0.6079 | -0.0167 |
| NMI | Bass | 0.3388 | -0.0092 |
| NMI | Bayespace | 0.00319 | 0.1058 |
| NMI | CCST | 1.14E-06 | 0.1967 |
| NMI | GraphPCA | 0.6462 | 0.0083 |
| NMI | Leiden | 3.01E-10 | 0.3750 |
| NMI | Louvain | 2.74E-09 | 0.3692 |
| NMI | NichePCA | 2.01E-08 | 0.1550 |
| NMI | STAGATE | 0.3028 | 0.0183 |
| NMI | Spaceflow | 5.70E-08 | 0.2800 |
| NMI | SpatialPCA | 0.8921 | -0.0025 |

**Table S3.** Results of the Wilcoxon signed-rank test assessing pairwise differences between jsPCA and benchmark methods based on ARI and NMI, using the DLPFC dataset.

| Metric | Method | p-value | Median diff |
| --- | --- | --- | --- |
| ARI | Bass | 0.0791 | 0.0200 |
| ARI | Bayespace | 0.0566 | 0.1250 |
| ARI | CCST | 0.00098 | 0.2150 |
| ARI | GraphPCA | 0.1240 | 0.0500 |
| ARI | Leiden | 0.00049 | 0.3800 |
| ARI | Louvain | 0.00049 | 0.3750 |
| ARI | NichePCA | 0.00049 | 0.3150 |
| ARI | STAGATE | 0.0322 | 0.0500 |
| ARI | Spaceflow | 0.00049 | 0.3600 |
| ARI | SpatialPCA | 0.5088 | -0.0200 |
| NMI | Bass | 0.1719 | 0.0200 |
| NMI | Bayespace | 0.00098 | 0.1300 |
| NMI | CCST | 0.00049 | 0.2150 |
| NMI | GraphPCA | 0.7109 | 0.0350 |
| NMI | Leiden | 0.00049 | 0.4200 |
| NMI | Louvain | 0.00049 | 0.4150 |
| NMI | NichePCA | 0.00049 | 0.1800 |
| NMI | STAGATE | 0.4502 | 0.0400 |
| NMI | Spaceflow | 0.00049 | 0.3150 |
| NMI | SpatialPCA | 0.5566 | 0.0050 |

**Table S4.** Results of the t-test assessing pairwise differences between jsPCA and benchmark methods based on ARI and NMI, using the first 14 sagittal sections of the MOSTA dataset.

| Metric | Method | p-value | Mean diff |
| --- | --- | --- | --- |
| ARI | Bass | 0.0846 | 0.0229 |
| ARI | GraphPCA | 0.000436 | 0.0529 |
| ARI | STAGATE | 0.000220 | 0.0500 |
| ARI | SpatialPCA | 9.96E-06 | 0.0936 |
| NMI | Bass | 0.0344 | 0.0214 |
| NMI | GraphPCA | 0.000165 | 0.0336 |
| NMI | STAGATE | 0.000122 | 0.0307 |
| NMI | SpatialPCA | 0.000119 | 0.0479 |

**Table S5.** Results of the Wilcoxon signed-rank test assessing pairwise differences between jsPCA and benchmark methods based on ARI and NMI, using the first 14 sagittal sections of the MOSTA dataset.

| Metric | Method | p-value | Median diff |
| --- | --- | --- | --- |
| ARI | Bass | 0.0784 | 0.025 |
| ARI | GraphPCA | 0.00185 | 0.040 |
| ARI | STAGATE | 0.00232 | 0.050 |
| ARI | SpatialPCA | 0.000979 | 0.085 |
| NMI | Bass | 0.0248 | 0.025 |
| NMI | GraphPCA | 0.00160 | 0.050 |
| NMI | STAGATE | 0.00177 | 0.035 |
| NMI | SpatialPCA | 0.00220 | 0.060 |

**Table S6.** Results of the t-test assessing pairwise differences between jsPCA and the benchmark methods (BASS and STAGATE), based on ARI and NMI, using the whole MOSTA dataset.

| Metric | Method | p-value | Mean diff |
| --- | --- | --- | --- |
| ARI | Bass | 1.72E-06 | 0.0283 |
| ARI | STAGATE | 2.91E-14 | 0.0558 |
| NMI | Bass | 4.12E-14 | 0.0402 |
| NMI | STAGATE | 1.63E-14 | 0.0290 |

**Table S7.** Results of the Wilcoxon signed-rank test assessing pairwise differences between jsPCA and the benchmark methods (BASS and STAGATE), based on ARI and NMI, using the whole MOSTA dataset.

| Metric | Method | p-value | Median diff |
| --- | --- | --- | --- |
| ARI | Bass | 1.31E-05 | 0.020 |
| ARI | STAGATE | 1.29E-09 | 0.050 |
| NMI | Bass | 6.25E-09 | 0.040 |
| NMI | STAGATE | 2.59E-09 | 0.040 |

**Table S8.** Ranking of marker genes across jsPCA components, for slice 151673 of the DLPFC dataset.

| Gene | jsPCA1 | jsPCA2 | jsPCA3 | jsPCA4 | jsPCA5 | jsPCA6 | jsPCA7 | jsPCA8 | jsPCA9 | jsPCA10 |
| --- | --- | --- | --- | --- | --- | --- | --- | --- | --- | --- |
| ADCYAP1 | 2105 | 257 | 486 | 6377 | 823 | 447 | 64 | 13472 | 4032 | 4091 |
| AQP4 | 236 | 515 | 612 | 8736 | 11 | 9629 | 5499 | 1631 | 131 | 26 |
| BCL11B | 7998 | 13409 | 833 | 8959 | 8110 | 3826 | 8458 | 6793 | 7908 | 14586 |
| CUX2 | 1359 | 38 | 232 | 970 | 201 | 14979 | 12411 | 7575 | 9893 | 1027 |
| HBB | 3042 | 13067 | 1637 | 4625 | 105 | 153 | 155 | 7168 | 726 | 27 |
| HPCAL1 | 1710 | 2 | 3026 | 35 | 1101 | 3239 | 5634 | 184 | 1030 | 3042 |
| IGKC | 3372 | 12218 | 61 | 612 | 43 | 466 | 845 | 1 | 4 | 1119 |
| KRT17 | 2341 | 327 | 14 | 24 | 27 | 7929 | 5 | 118 | 9886 | 8141 |
| MBP | 1 | 671 | 791 | 223 | 1836 | 4592 | 12547 | 10490 | 1205 | 6622 |
| MOBP | 6 | 1755 | 573 | 754 | 290 | 6192 | 13433 | 1409 | 13902 | 5698 |
| NPY | 1358 | 11356 | 4071 | 6842 | 10682 | 1279 | 3379 | 4450 | 9401 | 994 |
| NTNG2 | 10624 | 417 | 93 | 380 | 5401 | 8230 | 853 | 4539 | 12641 | 8994 |
| PCP4 | 580 | 11 | 7211 | 13830 | 527 | 10 | 1 | 147 | 204 | 48 |
| RELN | 5898 | 92 | 13067 | 1299 | 80 | 607 | 30 | 709 | 3643 | 4684 |
| RORB | 1233 | 88 | 599 | 309 | 9689 | 11390 | 12 | 3831 | 6925 | 12476 |
| SNAP25 | 80 | 9 | 580 | 155 | 13517 | 3354 | 1229 | 5272 | 1989 | 9720 |
| TRABD2A | 4689 | 91 | 2341 | 9183 | 8598 | 78 | 7 | 11778 | 4915 | 95 |

